# Membrane Anisotropy Reshapes Scale-Free Correlations and Directional Mechanical Susceptibility in Transmembrane Proteins

**DOI:** 10.64898/2026.08.20.745933

**Authors:** Jian Wang, Zizheng He, Xiaoyi Chen, Guannan Wang, Qian-Yuan Tang

**Affiliations:** College of Civil Engineering and Architecture, Zhejiang University, Hangzhou, 310058, China; Department of Physics, Hong Kong Baptist University, Kowloon Tong, Hong Kong SAR, China; Beijing Normal-Hong Kong Baptist University, Guangdong Provincial Key Laboratory of Interdisciplinary Research and Application for Data Science, Zhuhai, 519087, China

## Abstract

Long-range correlated motions couple distant regions of a protein, providing a physical basis for allosteric communication, cooperative conformational change, and the balance between structural stability and sensitivity to perturbations. Yet membrane proteins operate within a strongly anisotropic lipid bilayer, and how this environment reshapes such system-spanning coordination remains unclear. Using an implicit-membrane anisotropic network model, we perform a proteome-wide analysis of more than 3,000 human transmembrane proteins. We find that long-range correlations remain scale free under membrane constraints but become strongly direction dependent. Across protein sizes and topologies, their correlation lengths continue to scale with the corresponding molecular dimensions, while increasing membrane anisotropy extends in-plane correlations and shortens those along the membrane normal. Because spontaneous correlations and perturbation responses arise from the same underlying mechanics, we further resolve residue-level responses into in-plane and normal components. The resulting directional mechanical susceptibility provides new predictions of mutation-sensitive sites in GPCRs beyond those captured by conventional scalar flexibility measures. Together, these results show how environmental symmetry breaking can organize protein mechanics across scales, linking collective dynamics to the functional sensitivity of individual residues and connecting a general physical mechanism to experimentally measurable protein function.

## I. INTRODUCTION

Long-range correlated motions provide a physical basis for communication across proteins, coupling residues separated across the molecular structure and enabling distant regions to respond collectively [1–5]. Such system-spanning coordination provides a mechanism for balancing structural stability with sensitivity to local perturbations. The associated correlation length scales with molecular size rather than a fixed microscopic length, revealing a scale-free organization of protein dynamics analogous to collective behavior observed in other biological and soft-matter systems [6–13]. A fundamental question is whether this scale-free organization remains robust when the mechanical environment breaks spatial symmetry and constrains motion differently along different directions. Membrane proteins provide a natural setting for addressing this question: the lipid bilayer distinguishes the membrane plane from its normal and imposes direction-dependent mechanical constraints that influence helix tilting, lateral repacking, pore opening, signaling, and transport [14–20]. Yet previous studies have largely focused on individual proteins or specific functional processes, leaving unresolved whether long-range scale-free correlations persist under membrane confinement and how membrane-induced symmetry breaking reorganizes them between the membrane plane and its normal.

A second question follows directly from this directional collective picture: how does membrane anisotropy affect the mechanical sensitivity of individual residues? In elastic-network and linear-response descriptions of protein dynamics [21–25], equilibrium correlations and responses to local perturbations are governed by the same mechanical Hessian. Perturbation–response scanning (PRS) measures how a localized force propagates through this mechanical network [26–28], and the dynamic flexibility index (DFI) summarizes the resulting response as a residue-level susceptibility [29]. Such mechanical signatures have been associated with functional, evolutionary, and disease-relevant sites [30–38]. Standard DFI, however, treats this response as a scalar and therefore discards the distinction between motion in the membrane plane and along its normal. Direction-resolved PRS preserves this distinction [39], raising the question of whether membrane anisotropy creates residue-level mechanical signatures that are hidden in scalar measures. G protein-coupled receptors (GPCRs) provide a natural test because their activation requires coordinated rearrangements from the extracellular ligand-binding region, across the transmembrane bundle, to the intracellular signaling interface [40–48], while extensive mutational data allow this directional mechanical information to be tested against functionally sensitive sites [49–53].

In this paper, we employ the implicit-membrane anisotropic network model (imANM), which introduces direction-dependent stiffness while preserving native co-ordinates and contact topology [14], to investigate how membrane anisotropy organizes transmembrane protein (TMP) mechanics across scales. Across the human trans-membrane proteome, we show that long-range correlations retain their scale-free organization under membrane confinement but become strongly direction dependent: their characteristic lengths remain coupled to protein dimensions, while increasing anisotropy extends inplane correlations and shortens those along the membrane normal. We then resolve perturbation responses into the same two directional components to determine how this anisotropic collective mechanics is expressed at the level of individual residues. In GPCRs, the resulting directional susceptibility reveals complementary information about experimentally mutation-sensitive sites beyond conventional scalar flexibility measures. Together, these analyses establish a direct mechanical link between the directional reorganization of collective correlations and residue-level functional susceptibility.

## II. DIRECTIONAL ORGANIZATION OF SCALE-FREE CORRELATIONS IN TRANSMEMBRANE PROTEINS

### Anisotropic elastic network and directional correlations

To examine how membrane confinement reshapes long-range correlated motion, we employ the implicit-membrane anisotropic network model (imANM) [14]. As shown in Fig. 1a, the protein is represented by *N* nodes at their native *C*_*α*_ positions 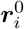, with native contacts connected within a cutoff *r*_*c*_ = 10 Å. Membrane anisotropy is introduced through direction-dependent stiffnesses while leaving the native coordinates and contact topology unchanged. For a native contact *r*_*ij*_ *< r*_*c*_, the off-diagonal Hessian block is

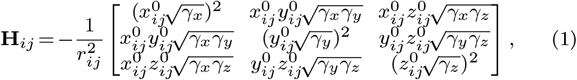

where 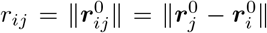, and the diagonal block **H**_*ii*_ *=* − ∑_*j*≠*i*_ **H**_*ij*_. With the membrane normal aligned with the *z* axis, we set *γ*_*x*_ = *γ*_*y*_ = *sγ*_*z*_, where *s* is the stiffness ratio [14]. Increasing *s* therefore strengthens the lateral mechanical constraint relative to the membrane normal uniformly across the native contact network.

**FIG. 1.**
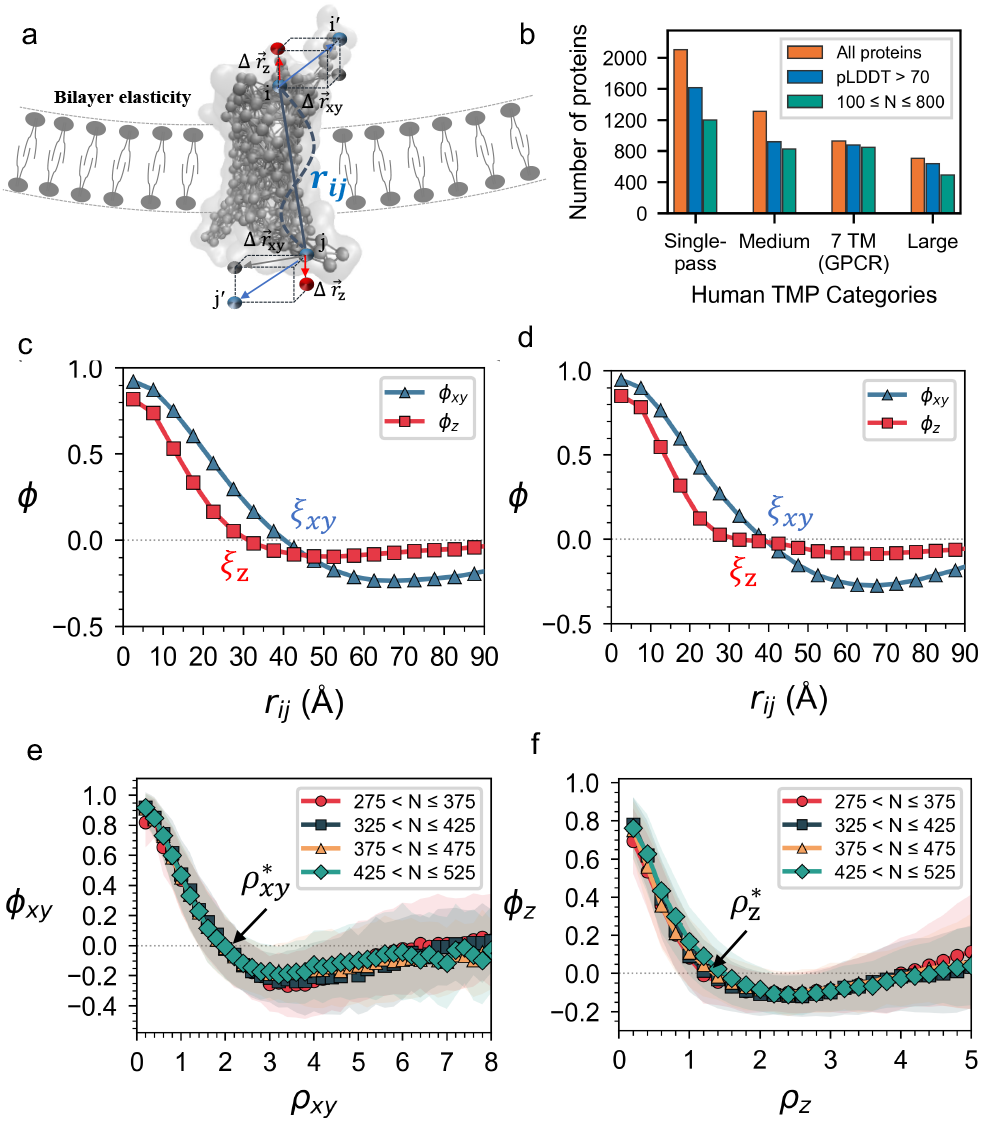
Directional decomposition of the correlated motions in transmembrane proteins. (a) Schematic of the imANM directional decomposition. (b) Dataset construction from TmAlphaFold. (c,d) Average directional correlation functions *ϕ*_*xy*_ and *ϕ*_*z*_ as functions of the three-dimensional native inter-residue distance *r*_*ij*_ at *s* = 16 for length-matched human TMPs (375–475 aa), including all four topological classes (a total of *n* = 552 proteins, panel c) and the 7TM subclass alone (a total of *n* = 120 proteins, panel d). (e,f) Rescaling the distance by the corresponding molecular extent collapses the average directional correlation functions across four protein-length bins (275–525 aa; a total of *n* = 1, 971 proteins), from which the reduced correlation lengths 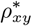 and 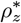 are obtained.

The collective fluctuations are determined by the 3*N* × 3*N* anisotropic Hessian **H**_*s*_, formed as an *N* × *N* block matrix with each block **H**_*ij*_ of size 3 × 3. After excluding the rigid-body zero modes, the equilibrium displacement covariance is proportional to the Moore–Penrose pseudoinverse, 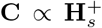, or equivalently reconstructed from the nonzero normal modes weighted by their inverse eigenvalues. Long-range correlations are dominated by the low-frequency collective spectrum, and the extracted correlation lengths are robust to the number of modes retained (Supp. Mat. Sec. 2). Each 3 × 3 covariance block describes the correlated displacements of a residue pair. We project these correlations onto the membrane plane and its normal and normalize by the corresponding residue fluctuations to obtain 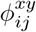 and 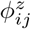. To characterize how these correlations depend on spatial separation, we follow the standard definition of distance-dependent correlation functions [6, 7, 11, 12] and construct *ϕ*_*xy*_(*r*) and *ϕ*_*z*_(*r*) by averaging the corresponding pairwise correlations over residue pairs at mutual distance *r*_*ij*_ in the native-state structure. Comparing *ϕ*_*xy*_(*r*) and *ϕ*_*z*_(*r*) therefore allows us to determine whether the two directional correlations show similar distance dependence or are reorganized differently by membrane anisotropy. Further computational details and robustness analyses are provided in Supp. Mat.

### Proteome-wide finite-size scaling of directional long-range correlations

With these directional correlation functions defined, we examined their behavior across transmembrane proteins (TMPs) of different sizes and topologies. We assembled *n*_p_ = 3359 human TMPs with mean pLDDT *>* 70 from the membrane-oriented AlphaFold structures in TmAlphaFold [54], which provides the membrane orientation with the membrane plane denoted as *xy* and its normal aligned along the *z* axis. The main analysis focuses on monomeric structures to isolate the effect of membrane anisotropy from additional inter-subunit interactions. We classified the proteins by the number of membrane-spanning transmembrane (TM) segments: single-pass (TM = 1), medium multi-pass (TM = 2–6), 7-pass (TM = 7, hereafter 7TM; predominantly GPCRs), and large multi-pass (TM *>* 7) (Fig. 1b). For each class, we analyzed the distance-dependent correlation functions *ϕ*_*xy*_(*r*) and *ϕ*_*z*_(*r*). Following previous finite-size correlation analyses [6, 11, 12], we define the directional correlation length *ξ*_*α*_ (*α*∈ { *xy, z}*) as the first zero of *ϕ*_*α*_(*r*). After removal of the rigid-body zero modes, positive correlations in a finite system are balanced by anticorrelations, so the first zero marks the distance over which positive correlations persist.

Consistent with previous imANM studies [14], we take *s* = 16 as a representative anisotropic condition; the main directional trends persist over a broad range of *s*. For the length-matched ensemble containing all four topological classes, with 375 *< N* ≤ 475 residues (a total of *n* = 552 proteins), the ensemble-averaged directional correlation functions separate clearly, with *ξ*_*xy*_ = 40.3 Å and *ξ*_*z*_ = 31.3 Å compared with a mean radius of gyration ⟨*R*_*g*_⟩ = 34.1 Å (Fig. 1c). The same pattern is observed within the 7TM subclass over the same length range (a total of *n* = 120 proteins), where *ξ*_*xy*_ = 38.9 Å, *ξ*_*z*_ = 32.2 Å, and ⟨*R*_*g*_⟩ = 33.2 Å (Fig. 1d). In both ensembles, *ξ*_*xy*_ *>* ⟨*R*_*g*_⟩ *> ξ*_*z*_, showing that the correlation length remains tied to the overall protein scale but is split directionally by membrane anisotropy, with more extended in-plane coordination and shorter normal correlations. The agreement between the pooled and 7TM ensembles further shows that this separation is not caused by mixing different transmembrane architectures. The in-plane correlation function also develops a pronounced negative tail at larger distances, indicating coordinated counter-motions between distant structural regions. The extracted correlation lengths remain nearly unchanged over a broad range of mode truncations and converge as the contact cutoff is increased, confirming that the directional separation is robust to these parameter choices (Fig. S2). For comparison, a representative dimeric assembly in the Supp. Mat. shows a similar separation between the *xy* and *z* correlation functions (Fig. S4) [55], indicating that the directional effect is not specific to the monomeric representation used in the main analysis.

The preceding analysis focused on proteins of comparable size. We next examined whether the directional organization of correlations persists across proteins of different sizes. In an approximately isotropic system, a single characteristic dimension such as the radius of gyration *R*_*g*_ can be used to rescale spatial distances. Here, however, the membrane distinguishes the in-plane and normal directions, so the relevant molecular size is itself direction dependent. We therefore characterize the molecular dimensions by the root-mean-square coordinate extents *L*_*xy*_ within the membrane plane and *L*_*z*_ along the membrane normal. Their ratio *η* = *L*_*z*_*/L*_*xy*_ provides a membrane-referenced measure of molecular anisotropy and shows trends consistent with the conventional gyration-tensor descriptors *b*^′^ and *κ*^2^ across all topological classes (Supp. Mat. Fig. S1).

Using these directional molecular extents, we next tested whether the in-plane and normal correlations retain a common finite-size organization as protein size increases. If so, their characteristic lengths should scale with the corresponding molecular dimensions rather than saturate at a fixed microscopic scale. We therefore rescale the three-dimensional inter-residue distance by *L*_*xy*_ for *ϕ*_*xy*_(*r*) and by *L*_*z*_ for *ϕ*_*z*_(*r*), and define the reduced correlation length 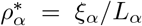 for *α* ∈ {*xy, z*}. Across four chain-length bins spanning 275–525 residues (a total of *n* = 1, 971 proteins), the correlation functions collapse onto separate in-plane and normal master curves (Fig. 1e,f), while 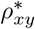 and 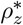 remain approximately independent of protein size. A similar directional collapse is observed across the four topological classes within the length-matched subset (375–475 residues, a total of *n* = 552 proteins; Supp. Mat. Fig. S2). These results show that membrane confinement does not truncate collective correlations to a fixed local scale: the correlation lengths remain coupled to protein dimensions, consistent with scale-free, critical-like organization. The membrane therefore preserves long-range collective coordination while breaking its directional symmetry, organizing correlations differently within the membrane plane and along its normal.

### Directional correlation lengths under membrane anisotropy

The preceding analysis established the directional correlation behavior from ensemble-averaged profiles. We next tested whether this reorganization persists at the level of individual proteins by varying the stiffness ratio over 1 ≤ *s* ≤ 64 and examining the protein-specific reduced correlation lengths, focusing first on proteins with similar chain lengths (375–475 residues). Across all topological classes, 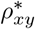 increases with *s*, whereas 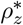 decreases (Fig. 2a,b). This opposing response varies across topologies and is most evident in the 7TM class. For 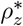, the decrease becomes relatively modest beyond *s* = 16, whereas 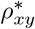 continues to increase. At *s* = 16, the two directional correlation lengths are already well separated; this value also coincides with the representative anisotropic condition used in previous imANM studies [14].

**FIG. 2.**
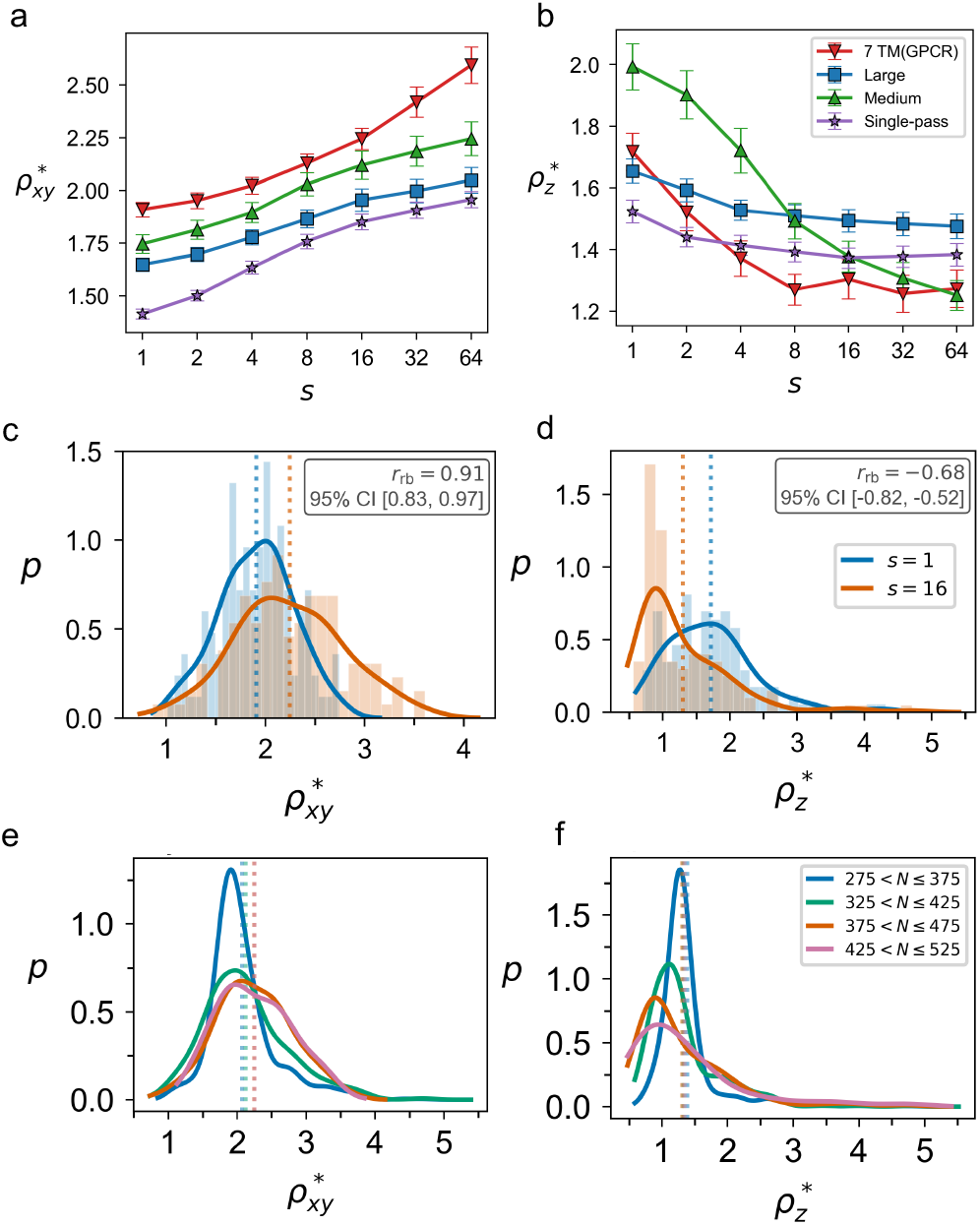
Opposite changes in directional correlation lengths under membrane anisotropy. (a,b) Reduced directional correlation lengths 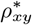 and 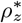 as functions of the stiffness ratio *s* for length-matched transmembrane proteins (375–475 residues) across four topological classes. (c,d) Probability densities of 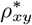 and 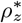 for the 7TM subclass over the same length range (a total of *n* = 120 proteins), comparing *s* = 1 and *s* = 16. (e,f) Probability densities of 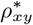 and 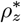 for 7TM proteins across four chain-length bins spanning 275–525 residues (a total of *n* = 778 proteins) at *s* = 16.

The contrast between the isotropic condition (*s* = 1) and the representative anisotropic condition (*s* = 16) can be quantified at the level of individual proteins. Within the 7TM subset (375–475 residues, a total of *n* = 120 proteins), the mean reduced in-plane correlation length increases from 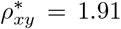 at *s* = 1 to 2.24 at *s* = 16, whereas the mean reduced normal correlation length decreases from 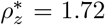 to 1.30 (Fig. 2c,d). With paired differences defined as *s* = 16 minus *s* = 1, matched-pairs rank-biserial correlations give *r*_rb_ = 0.91 for the in-plane comparison and *r*_rb_ = − 0.68 for the normal comparison, with confidence intervals estimated by percentile bootstrap using 5,000 resamples. These oppositely signed effect sizes show that the directional shifts occur consistently across individual proteins rather than reflecting only a change in the ensemble average. Similar responses are observed in the other topological classes (Fig. S3c,d).

To determine whether this directional response persists across protein sizes, we examined 7TM proteins spanning 275–525 residues (a total of *n* = 778 proteins). At *s* = 16, both 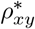 and 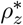 remain nearly unchanged across four chain-length bins (Fig. 2e,f), with similar behavior in the other topological classes (Fig. S3e,f). Thus, the opposing changes in in-plane and normal correlations are not a consequence of protein size, but represent a persistent feature of membrane-constrained mechanics across molecular scales. Even as TMPs become larger, membrane anisotropy continues to reorganize their system-spanning correlations in the same directional manner, while topology mainly modulates the magnitude of this response.

### Membrane constraint reshapes residue-level correlation patterns

The opposing shifts in the reduced correlation lengths suggest that membrane anisotropy reorganizes the underlying residue-level correlation pattern. To visualize how this reorganization occurs at the level of individual residue pairs, we examined the full direction-resolved correlation matrices as the stiffness ratio *s* increased, using human cytochrome *c* oxidase subunit 2 (P00403, *N* = 227) as an illustrative example. As shown in Fig. 3, the *xy* correlation pattern at low *s* consists of relatively small, distinct blocks. With increasing *s*, these blocks broaden and connect across more distant sequence regions, forming larger coherent domains, accompanied by an increase in the mean correlation between contacting residues. The *z* direction shows the opposite trend: correlations spanning extended regions progressively weaken and fragment into more localized patches, together with a decrease in the mean contact correlation. These residue-level changes provide a direct microscopic picture of the preceding correlation-length analysis: the growth and merging of in-plane correlated domains allow positive correlations to persist over longer distances, increasing 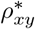, whereas the fragmentation of normal correlations shortens their persistence and decreases 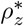. A similar contact-level signature is observed in dimeric assemblies, where increasing anisotropy increases the mean contact correlation in the *xy* direction while decreasing it along *z* (Fig. S4). This consistency indicates that the directional reorganization is not specific to the monomeric representation used in the main analysis.

**FIG. 3.**
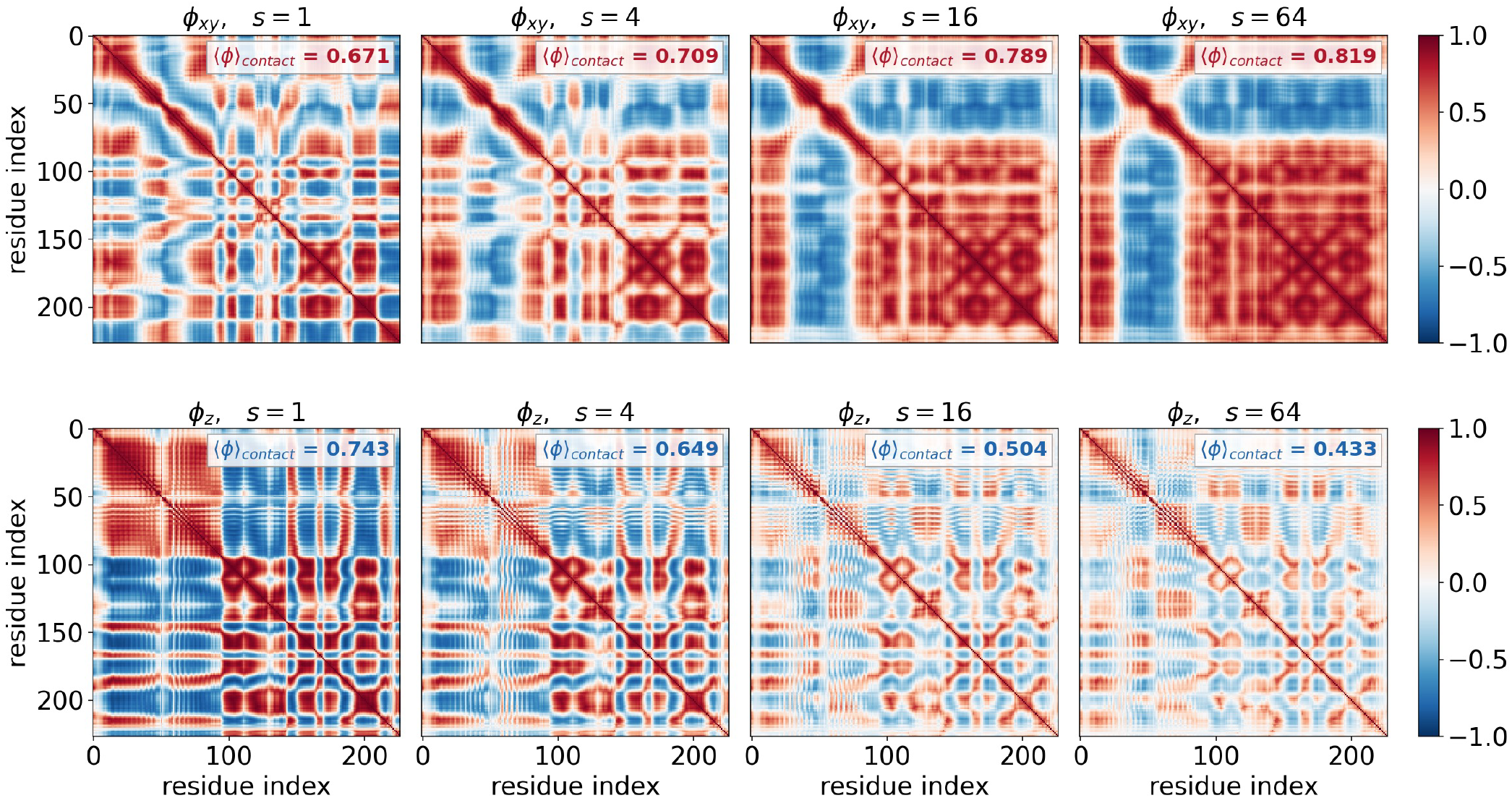
Membrane constraint reshapes residue-level correlation patterns. Direction-resolved correlation matrices for human cytochrome *c* oxidase subunit 2 (P00403, *N* = 227), used here as an illustrative example, showing 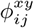 (top row) and 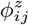 (bottom row) at stiffness ratios *s* = 1, 4, 16, and 64. Residues are ordered sequentially. The mean directional correlation 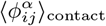 over residue pairs in contact within *r*_*ij*_ *< r*_*c*_ = 10 Å is also shown in each panel.

This opposing reorganization follows from the mechanical compatibility of the native contact network. Because native contacts are generally tilted relative to the membrane, extended normal motion requires coordinated lateral rearrangements to accommodate the contact geometry. Increasing the lateral constraint therefore has two complementary consequences. Within the membrane plane, relative motion between neighboring regions is suppressed, effectively locking smaller dynamical units into larger coherent domains. Along the membrane normal, the same locking restricts the distributed lateral accommodation required to coordinate distant regions, causing long-range correlations to break into localized patches. The contact-level compatibility condition and the corresponding spectral description of this mechanism are derived in Supp. Mat. Sec. 3.

## III. DIRECTIONAL RESPONSES REVEAL MUTATION-SENSITIVE SITES

The preceding analysis shows that membrane anisotropy reorganizes long-range correlations between the membrane plane and its normal while preserving their system-spanning character. A natural next question is whether this directional organization is also expressed in the mechanical susceptibility of individual residues. Because spontaneous correlations and force-induced responses arise from the same underlying mechanics, directional perturbation responses provide a direct link between the collective organization established above and residue-level sensitivity. GPCRs provide a particularly suitable test because their function requires coordinated rearrangements across the TM bundle and extensive experimental mutagenesis data are available in GPCRdb [56–58]. We therefore ask whether membrane-resolved mechanical susceptibility provides information about mutation-sensitive sites beyond that contained in scalar mechanical measures.

### Direction-resolved perturbation–response analysis

We combine imANM with perturbation–response scanning (PRS) [26, 27] to examine how the same anisotropic mechanics that governs equilibrium correlations is expressed in residue-level responses. By resolving the response into in-plane and normal components, we extend the directional analysis of correlated fluctuations to mechanical susceptibility and define direction-resolved DFI [29]. As illustrated in Fig. 4a, PRS applies a fictitious unit force ***F*** to each residue *j*, sampled along seven representative unoriented axes: the three Cartesian axes and four body diagonals of a cube. After excluding the zero modes, the resulting linear response is determined by the pseudoinverse of the anisotropic Hessian,

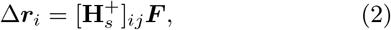

where 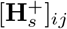 is the 3 × 3 block that maps a perturbation at residue *j* to the displacement of residue *i*.

**FIG. 4.**
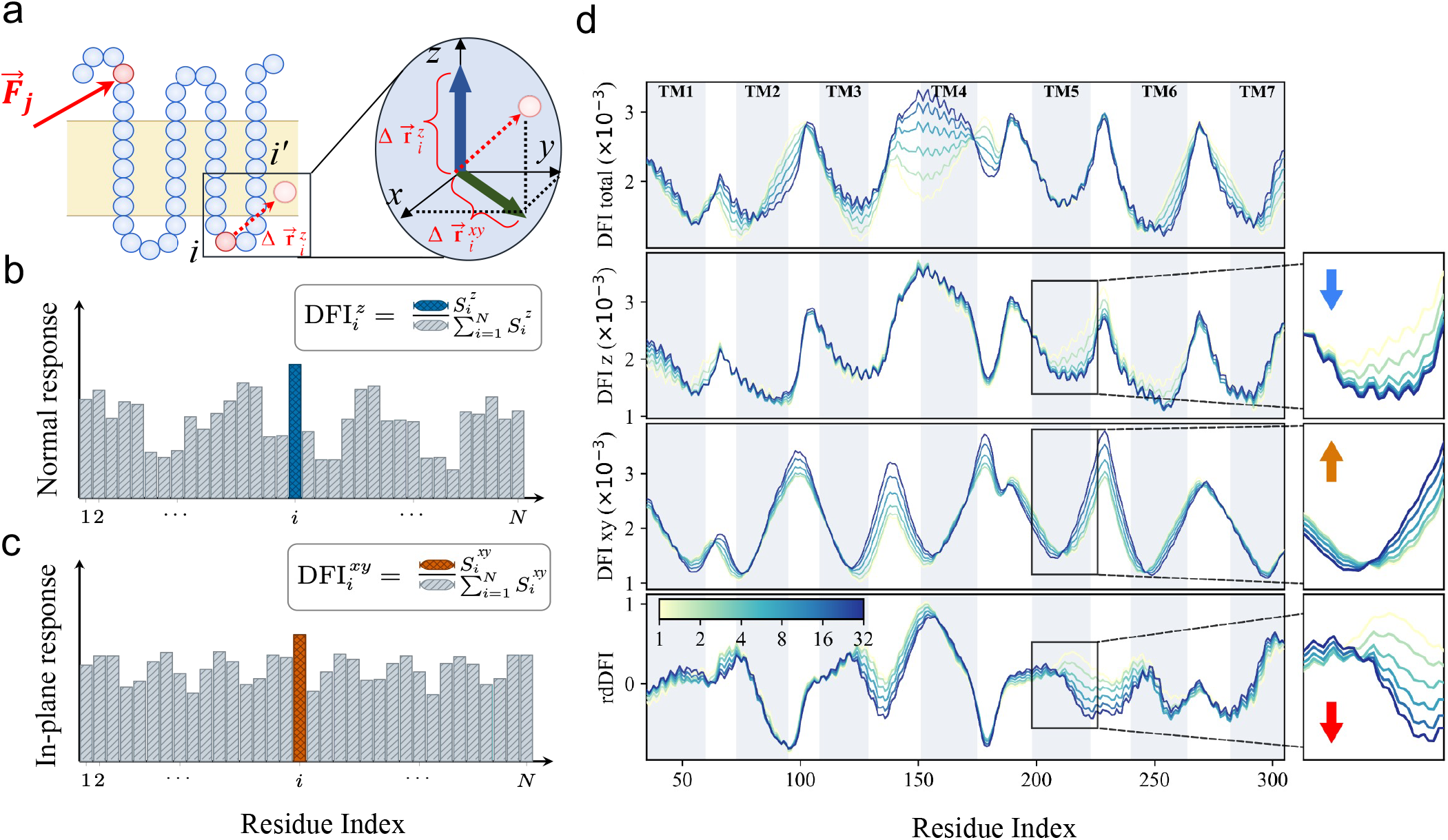
Directional decomposition of residue-level perturbation responses. (a–c) Schematics of direction-resolved PRS within the imANM framework and the construction of DFI^*z*^ and DFI^*xy*^ from the normal and in-plane response components. (d) Residue-level profiles of the direction-unresolved DFI, DFI^*xy*^, DFI^*z*^, and their log contrast rdDFI for the representative GPCR CCR1 at different stiffness ratios *s*. Enlarged views around TM5 highlight how the directional responses and the resulting relative directional DFI (rdDFI) contrast evolve with increasing *s*.

To retain the membrane-defined directional information, we project each displacement onto the membrane plane and its normal before taking its magnitude. With P_*xy*_ = diag(1, 1, 0) and P_*z*_ = diag(0, 0, 1), the force-orientation-averaged response matrix is

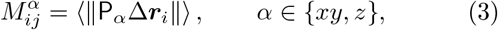

where ⟨· · · ⟩ denotes the average over force orientations. Summing over perturbation sites gives 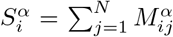. Normalizing these responses (Fig. 4b,c), gives

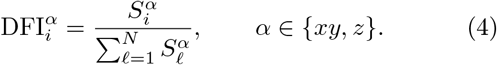

Thus, 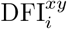 and 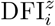 quantify the relative contribution of residue *i* to the in-plane and normal responses, respectively. Standard DFI uses the full three-dimensional response of the isotropic network, whereas imDFI applies the same direction-unresolved measure to imANM.

To quantify the directional imbalance of each residue, we introduce the relative directional DFI (rdDFI),

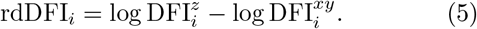

Because DFI^*z*^ and DFI^*xy*^ are normalized independently, rdDFI measures the directional bias of residue *i* relative to the overall response balance of the protein, rather than the absolute difference between the two response components. Positive rdDFI indicates relative enrichment of the normal response, whereas negative rdDFI indicates relative enrichment of the in-plane response.

As *s* increases, the directional response profiles progressively separate within the TM region of CCR1 (Fig. 4d). TM residues lose relative weight within the normal-response profile while gaining relative weight within the in-plane profile, consistent with the directional reorganization observed in the correlation analysis. The direction-unresolved response changes much less, indicating that this redistribution is largely hidden when the two components are combined. Because the contrast between DFI^*z*^ and DFI^*xy*^ varies strongly along the sequence, rdDFI further resolves how this directional bias differs from residue to residue. For rdDFI-based site identification, the profile is standardized within the TM region as *Z*_*i*_ = (rdDFI_*i*_ − *µ*_TM_)*/σ*_TM_, where *µ*_TM_ and *σ*_TM_ denote the mean and standard deviation, respectively, of the rdDFI values over all TM residues of the receptor. Residues with *Z*_*i*_ *<* − 1.0 therefore represent sites with strong relative enrichment of the in-plane response and are retained for comparison with experimental mutation data.

### Directional contrast identifies mutation-sensitive sites in GPCRs

We next tested whether directional mechanical susceptibility is associated with experimentally observed mutation sensitivity. We curated mutagenesis data from GPCRdb [56–58] for human non-olfactory receptors with reliable structural models. Both candidate-site identification and experimental benchmarking were restricted to residues within the TM helices, where membrane-defined anisotropic constraints are most directly relevant and the structural framework is comparatively conserved across GPCRs. The GPCRdb data include different experimental readouts, such as binding affinity, EC_50_, and IC_50_, with mutational effects reported as fold changes relative to the corresponding wild-type receptor. We classified a TM position as mutation sensitive if at least one experimentally tested mutation produced a 10-fold or greater increase or decrease; experimentally characterized positions with smaller effects were classified as non-sensitive. Because our analysis focuses on residue positions rather than specific amino-acid substitutions, multiple measurements at the same position were represented by the largest observed mutational effect relative to the wild type, irrespective of substitution, experimental condition, or readout. The benchmark therefore asks whether perturbing a given TM position can produce a strong functional effect.

To ensure a sufficient number of experimentally sensitive sites for receptor-level evaluation, we required each receptor to contain at least three mutation-sensitive TM positions. Among receptors satisfying this criterion, we selected the 20 with the largest numbers of experimentally characterized TM positions, thereby maximizing mutational coverage within this eligible receptor set. The resulting benchmark contained 1,452 unique experimentally characterized TM positions across the 20 receptors (Supplementary material). For each receptor, the hit rate was defined as the fraction of mechanically prioritized sites classified as mutation sensitive, considering only TM positions with available mutagenesis measurements.

Within this curated dataset, we compared three mechanical measures: isotropic DFI calculated at *s* = 1, direction-unresolved imDFI calculated from imANM at *s* = 16, and the directional contrast rdDFI at *s* = 16. This hierarchy distinguishes two effects of the membrane: changes in the overall mechanical response and redistribution of that response between the membrane plane and its normal. It therefore tests whether mutation sensitivity is associated primarily with total mechanical susceptibility or with its directional organization.

The sites prioritized by rdDFI overlap only weakly with those identified by isotropic DFI. For each mechanical measure, residue scores were standardized separately within the TM region, and the same cutoff, *Z <* − 1, was then used to define mechanically prioritized sites. Across the 20 receptors, this criterion identified 303 sites for DFI and 286 sites for rdDFI, of which only 62 were shared (Fig. 5a). The weak overlap therefore reflects differences in the underlying mechanical profiles rather than differences in the selection criterion. Directional contrast does not simply reorder residues along the same scalar flexibility axis; rather, it exposes sites with unexceptional total susceptibility but a pronounced bias toward one membrane-defined direction.

**FIG. 5.**
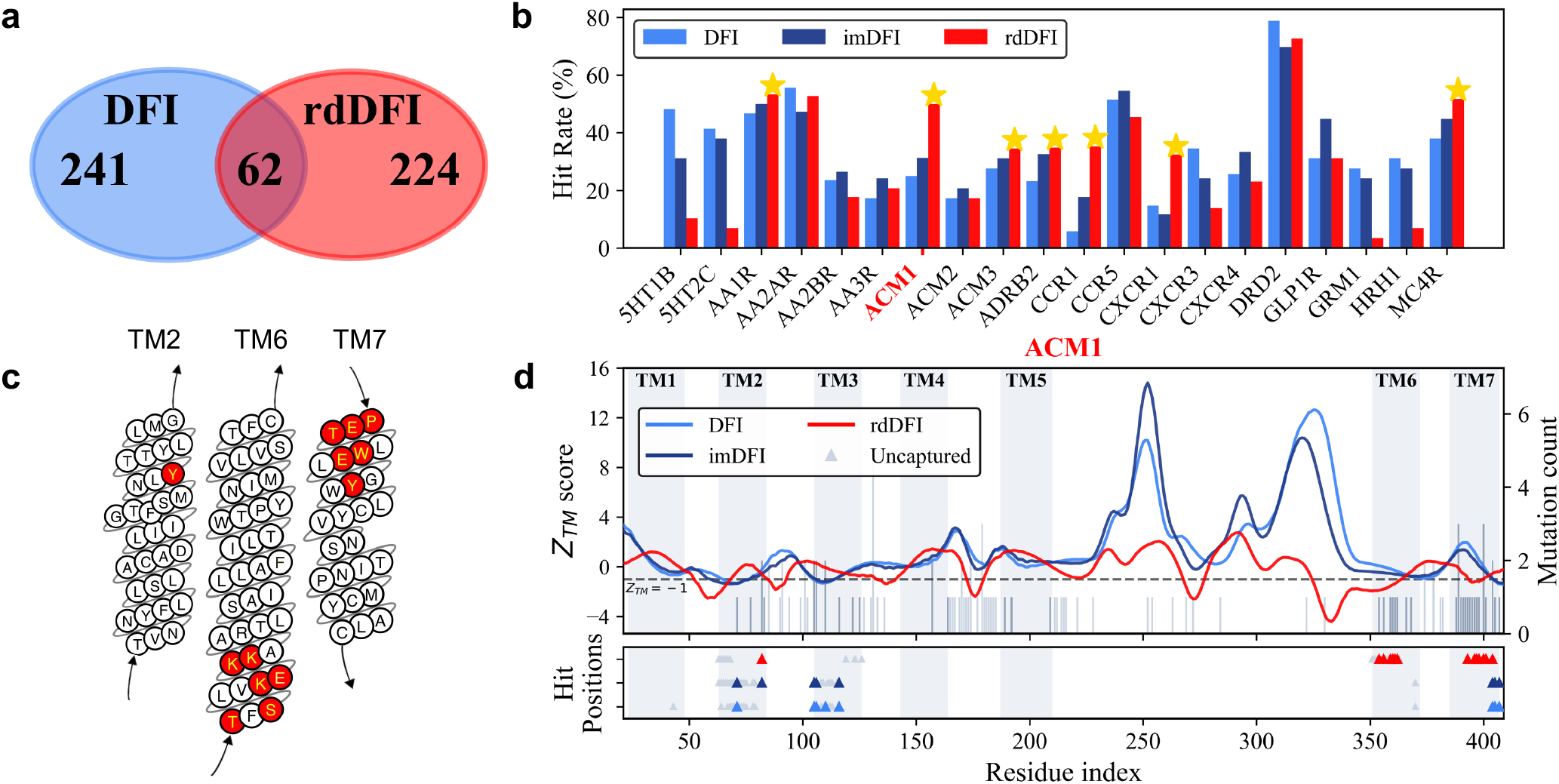
Relative directional DFI (rdDFI) reveals complementary mutation-sensitive sites in GPCRs. (a) Overlap between sites prioritized by DFI and rdDFI across 20 curated GPCRs. (b) Per-receptor hit rates of DFI, imDFI, and rdDFI against GPCRdb mutagenesis data; yellow stars indicate receptors for which rdDFI achieves the highest hit rate. (c) Experimentally mutation-sensitive sites prioritized by rdDFI in the muscarinic acetylcholine receptor M1 (ACM1), located on TM2, TM6, and TM7. The arrows indicate the N-to-C-terminal direction. (d) Sequence profiles of the experimental mutation data and the three mechanical measures for ACM1.

This directional information is also reflected in the experimental benchmark. Across the curated receptor set, rdDFI achieves hit rates comparable to those of isotropic DFI and imDFI for most receptors, while showing higher hit rates in several cases, with ACM1, CCR5, and CXCR3 providing particularly clear examples (Fig. 5b). Together with the limited overlap between prioritized sites, these results indicate that rdDFI provides complementary mechanical information beyond scalar flexibility. Notably, imDFI already incorporates membrane anisotropy into the mechanical network but combines the directional responses into a single scalar measure, and its residue-level profile remains very similar to isotropic DFI (Fig. 5d). Thus, introducing anisotropy alone adds relatively little information at the level of scalar susceptibility; the distinct signal emerges when the response is resolved into the membrane plane and its normal, indicating that the relevant information lies in the directional organization of mechanical susceptibility.

As a representative example, we examined the muscarinic acetylcholine receptor M1 (ACM1) in detail (Fig. 5c,d). The strongest rdDFI signals are concentrated in TM6 and TM7, where many experimentally mutation-sensitive positions are also found, whereas isotropic DFI and imDFI show much weaker distinction in these regions. The relevant mechanical feature is therefore not unusually low total flexibility, but how the allowed response is partitioned between the membrane plane and its normal. Conventional DFI preferentially highlights mechanically constrained sites in the orthosteric and structural core, including D105^3.32^, Y106^3.33^, C407^7.42^, and the conserved sodium-binding residue D71^2.50^ [59– 61]. Here and below, superscripts denote Ballesteros– Weinstein generic residue positions [62, 63]. In contrast, rdDFI highlights sites such as W400^7.35^ and Y82^2.61^, which have established roles in allosteric modulation and subtype-selective signaling [60, 64, 65]. Particularly notable is the strong relative enrichment of in-plane response toward the cytoplasmic end of TM6, the region that undergoes the characteristic outward rearrangement during receptor activation and forms part of the G-protein-coupling interface [41, 66].

The directional susceptibility revealed by rdDFI suggests that functionally important residues need not be characterized by uniformly low flexibility, but can instead be mechanically constrained in a direction-dependent manner. This provides a natural extension of the conventional hinge picture, in which low-DFI residues act as relatively rigid sites that organize collective motion [31, 32]. In membrane proteins, some residues may play a similar organizing role through directional rather than uniform constraint: their response is relatively suppressed along the membrane normal while remaining enriched within the membrane plane. We therefore refer to such sites as directional hinges, whose mechanical role is to restrict the directions available to collective rearrangements rather than simply resist motion. TM6 provides a representative example, where pronounced in-plane susceptibility is consistent with the lateral rearrangement required during receptor activation and G-protein coupling. In this sense, rdDFI adds information absent from scalar flexibility—it distinguishes not only how strongly a residue responds, but how the membrane organizes the geometry of that response.

## IV. DISCUSSION

In this study, we show that the lipid bilayer acts not simply as a mechanical constraint but as a symmetry-breaking environment that directionally reorganizes collective protein mechanics. Across the human transmembrane proteome, long-range correlations persist under membrane confinement, while their characteristic lengths separate between the membrane plane and its normal: increasing anisotropy extends in-plane coordination and shortens normal correlations, with both remaining coupled to protein size. The same directional organization appears at residue resolution, where membrane-resolved susceptibility provides complementary information about experimentally mutation-sensitive TM sites beyond conventional scalar flexibility measures.

The physical origin of this reorganization lies in how membrane anisotropy reshapes mechanical compatibility across the native contact network. Because tilted contacts couple lateral and normal displacements, increasing anisotropy changes which collective deformations can remain compatible across many contacts and thereby reorganizes the low-frequency collective response. The opposing shifts of the reduced directional correlation lengths are therefore two manifestations of the same redistribution of collective modes. Importantly, this reorganization does not destroy the underlying finite-size organization: the correlation lengths continue to grow with the corresponding molecular dimensions rather than approaching a fixed microscopic scale [6, 7, 67]. Membrane anisotropy thus changes the geometry of collective motion without eliminating its scale-free character.

At the residue level, this directional organization adds a distinct layer to existing perturbation–response descriptions of protein mechanics. Conventional DFI characterizes the overall mechanical susceptibility of a residue, while the Dynamic Coupling Index (DCI) measures preferential mechanical coupling between specific residues and functional sites [68]. rdDFI addresses a different aspect of the response by resolving how residue-level susceptibility is partitioned between the membrane plane and its normal. Residues with similar total flexibility can therefore exhibit markedly different directional susceptibilities and, consequently, different mechanical roles. This naturally extends the conventional mechanical-hinge picture: a functionally important site need not be uniformly rigid, but may instead constrain one component of motion while remaining responsive along another, thereby shaping the geometry of collective conformational change. Such directional hinges are particularly natural in membrane proteins, where the bilayer provides a physical reference frame for motion. The same directional decomposition could also be combined with DCI to resolve not only which residues are mechanically coupled, but also the directions along which this coupling is transmitted. Direction-resolved susceptibility therefore complements existing perturbation–response measures [49, 51], as well as deep mutational scanning, evolutionary constraints, and sequence- or structure-based variant-effect models [69–73], by providing a physical readout of how membrane geometry shapes site-specific functional sensitivity.

A limitation of the present framework is that it captures near-native directional mechanics rather than the full conformational landscape. It does not explicitly describe anharmonic transitions, contact rearrangements, or lipid- and state-specific interactions. TmAlphaFold provides the structural coverage required for proteome-wide analysis but not the conformational ensembles accessible through molecular dynamics or experiment. Likewise, the GPCRdb benchmark reflects the experimentally explored mutational landscape and is therefore shaped by the available mutagenesis coverage. Comparisons across functional states and more systematic mutational and conformational measurements will be important for determining how directional mechanical organization evolves with receptor conformation and contributes to function.

More broadly, the present results suggest a multiscale picture in which membrane geometry, molecular assembly, and local energetics jointly organize collective mechanics. Here, the membrane is represented as a uniform planar environment with a single global axis of anisotropy. In cellular membranes, variations in thickness, composition, tension, and curvature can make this directional constraint spatially heterogeneous. In curved regions such as cilia, membrane tubules, or mitochondrial cristae, the local membrane normal varies across space, suggesting that collective correlations may instead be organized by a spatially varying mechanical field. Molecular assembly may further propagate directional correlations across subunit interfaces [74], while local energetic and frustration landscapes determine where collective deformation is accommodated or concentrated [75]. Together, these ingredients point toward a broader framework in which membrane geometry defines directional constraints, molecular architecture governs their propagation, and local energetics shapes their expression as site-specific functional sensitivity. Similar principles may also guide the design of elastic networks capable of long-range mechanical communication [5, 76].

## Supporting information

This Supplementary Material provides supporting definitions, robustness tests, mechanistic analyses, oligomeric results, data, and code.

## DATA AVAILABILITY

All data and code supporting the findings of this study are available within the article and the Supplemental Material. Information on the source datasets, curated benchmark data, analysis code, and associated public repositories is provided in the Supplemental Material.

## ACKNOWLEDGMENTS

The authors thank Catherine H. H. Hor, Weitong Ren, Wenfei Li, Chak Leung Wong, and Tianlang Wu for helpful discussions. The authors acknowledge support from Natural Science Foundation of China (Nos. 12305052, 12322206, 12272055), Research Grants Council of Hong Kong (22302723), Guangdong Basic and Applied Basic Research Fund (2026A1515011563), Guangdong and Hong Kong Universities “1+1+1” Joint Research Collaboration Scheme (2025A0505000011), Hong Kong Baptist University’s funding support (RC-FNRA-IG/22-23/SCI/03, and RC-SFCRG/23-24/SCI/06).

