## Supplementary material for "Membrane Anisotropy Reshapes Scale-Free Correlations and Directional Mechanical Susceptibility in Transmembrane Proteins": This Supplementary Material provides supporting definitions, robustness tests, mechanistic analyses, oligomeric results, data, and code.

This Supplementary Material provides additional definitions and analyses supporting the main text, including directional molecular extents and correlation measures, robustness and finite-size scaling tests, the mechanical origin of the opposing directional reorganization, and an extension to oligomeric membrane-protein assemblies. In addition, the associated curated datasets and analysis code are provided as supplementary data and code repositories.

### S1. DIRECTIONAL MOLECULAR EXTENTS AND SCALE-FREE CORRELATION LENGTHS

#### S1.1 Directional molecular extents and geometric anisotropy

To characterize the size and shape of a transmembrane protein relative to the membrane, we define separate molecular extents within the membrane plane and along its normal. Let $\mathbf{r}_i = (x_i, y_i, z_i)$ denote the $C_\alpha$ coordinate of residue $i$, and let $\mathbf{r}_c = (x_c, y_c, z_c) = N^{-1}\sum_{i=1}^{N}\mathbf{r}_i$ be the geometric centroid.

The in-plane and normal molecular extents are defined as

$$L_{xy} = \left\{ \frac{1}{N}\sum_{i=1}^{N}\left[(x_i - x_c)^2 + (y_i - y_c)^2\right]\right\}^{1/2}. \tag{S1}$$

$$L_z = \left[\frac{1}{N}\sum_{i=1}^{N}(z_i - z_c)^2\right]^{1/2}. \tag{S2}$$

The two quantities give the characteristic molecular extents selected by the membrane. Their ratio,

$$\eta = \frac{L_z}{L_{xy}}, \tag{S3}$$

measures geometric anisotropy relative to the membrane. With the present definition, an isotropic three-dimensional structure has $L_{xy} = \sqrt{2}\,L_z$ and therefore $\eta = 1/\sqrt{2}$; deviations from this value reflect preferential extension within the membrane plane or along its normal. By construction, $R_g^2 = L_{xy}^2 + L_z^2$, so $L_{xy}$ and $L_z$ decompose the conventional overall molecular size into membrane-referenced components.

For comparison, let $\sigma_1 \geq \sigma_2 \geq \sigma_3$ denote the eigenvalues of the gyration tensor. The normalized asphericity $b' = [\sigma_1 - (\sigma_2+\sigma_3)/2]/(\sigma_1+\sigma_2+\sigma_3)$ measures the dominance of the largest principal dimension, whereas the relative shape anisotropy $\kappa^2 = 1 - 3(\sigma_1\sigma_2 + \sigma_2\sigma_3 + \sigma_3\sigma_1)/(\sigma_1+\sigma_2+\sigma_3)^2$ measures the overall departure from a spherical shape. The membrane-referenced ratio $\eta$ is strongly associated with both descriptors across all topological classes (Fig. S1).

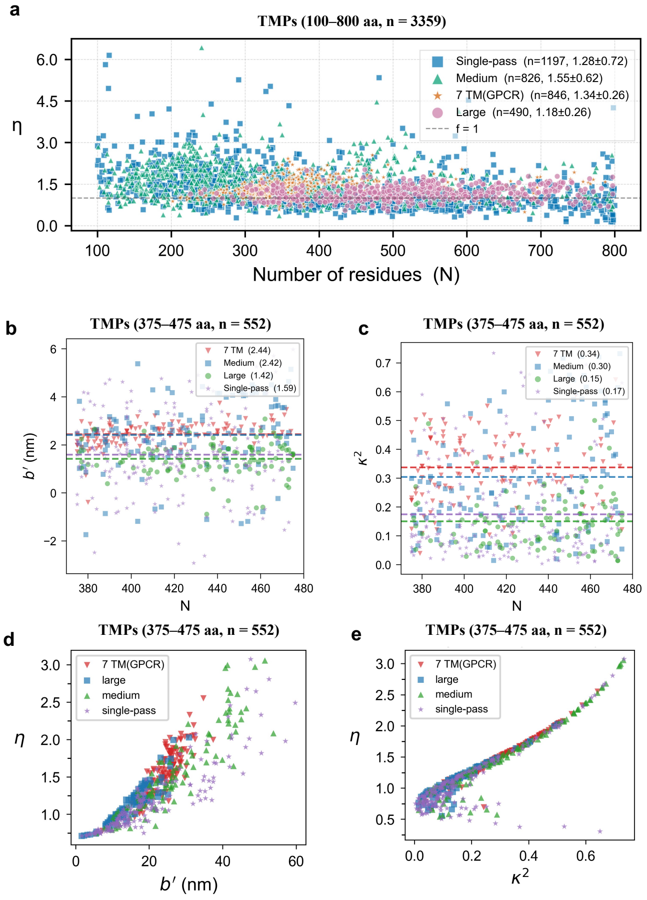


FIG. S1. Topology-dependent geometric anisotropy of transmembrane proteins. (a) Distribution of the directional extent ratio $\eta = L_z/L_{xy}$ over the full sequence-length range for single-pass, medium multi-pass, 7TM (predominantly GPCRs), and large multi-pass proteins. (b,c) Size-controlled comparison ($375 < N \leq 475$) of the conventional shape descriptors, asphericity $b'$ and relative shape anisotropy $\kappa^2$. (d,e) Relationships between $\eta$ and the gyration-tensor descriptors $b'$ and $\kappa^2$, respectively.

#### S1.2 Directional correlation functions

With the membrane normal aligned with the $z$ axis, we use the projectors $\mathsf{P}_{xy} = \mathrm{diag}(1,1,0)$ and $\mathsf{P}_z =$

diag(0, 0, 1). Then, the normalized directional correlation between residues $i$ and $j$ is defined as

$$\phi_{ij}^{\alpha} = \frac{\mathrm{Tr}(\mathsf{P}_\alpha C_{ij})}{\sqrt{\mathrm{Tr}(\mathsf{P}_\alpha C_{ii})\,\mathrm{Tr}(\mathsf{P}_\alpha C_{jj})}}, \qquad \alpha \in \{xy, z\}, \quad \text{(S4)}$$

where $C_{ij}$ is the $3 \times 3$ covariance block between residues $i$ and $j$.

For each protein, the direction-resolved correlation profile $\phi_\alpha(r)$ is obtained by binning $\phi_{ij}^{\alpha}$ according to the common three-dimensional Euclidean distance $r_{ij} = \|\boldsymbol{r}_{ij}^0\|$, using bins of width $\Delta r = 5$ Å:

$$\phi_\alpha(r) = \frac{1}{M(r)} \sum_{|r_{ij}-r| \le \Delta r/2} \phi_{ij}^{\alpha}, \qquad \alpha \in \{xy, z\}, \quad \text{(S5)}$$

where $M(r)$ is the number of residue pairs from that protein in the distance bin centered at $r$.

For ensemble analyses, these protein-specific profiles were evaluated on a shared distance grid with bin centers at $2.5, 7.5, 12.5, \ldots$ Å and then averaged bin-wise within each category. Each protein was assigned equal weight in this averaging, yielding the ensemble-averaged directional correlation curves.

Following the standard definition for distance-dependent correlations, the directional correlation length $\xi_\alpha$ is defined as the first zero of $\phi_\alpha(r)$:

$$\xi_\alpha = \min\{r > 0 : \phi_\alpha(r) = 0\}, \qquad \alpha \in \{xy, z\}. \quad \text{(S6)}$$

The zero is obtained by linear interpolation between adjacent bins bracketing the first sign change. Applied to a protein-specific profile, this yields the individual-protein correlation length; applied to the ensemble-averaged profile, it yields an ensemble-level correlation length, which is generally not equal to the mean of the individual-protein correlation lengths. For individual proteins, the reduced correlation lengths are defined as $\rho_{xy}^* = \xi_{xy}/L_{xy}$ and $\rho_z^* = \xi_z/L_z$.

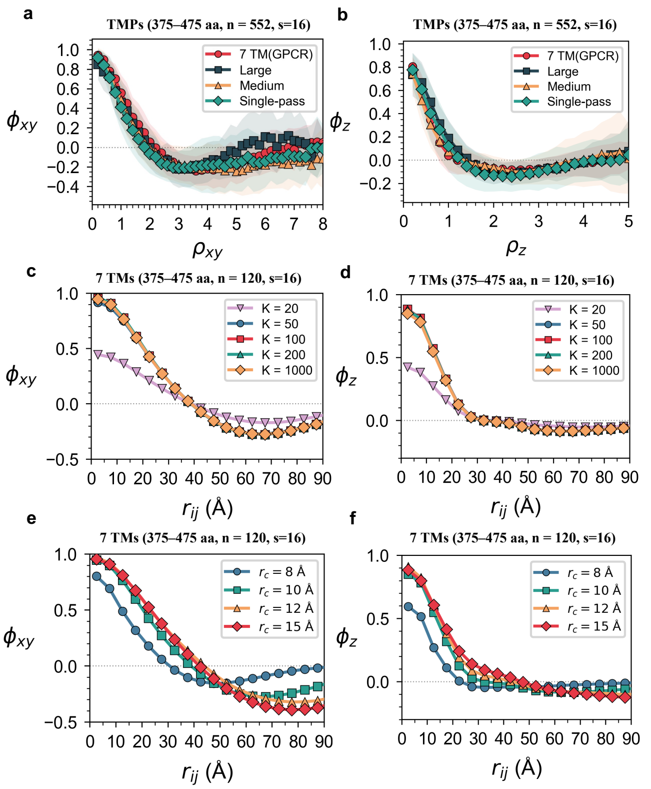


FIG. S2. Robustness and finite-size scaling of average directional correlations. (a,b) Size-normalized directional correlation functions for a length-matched ensemble of 552 transmembrane proteins ($375 < N \le 475$ residues) spanning four topological classes at $s = 16$. (c,d) Direction-resolved correlation functions $\phi_{xy}$ and $\phi_z$ as functions of the common three-dimensional inter-residue distance $r_{ij}$, reconstructed using progressively increasing numbers of nonzero modes ($K = 20, 50, 100, 200$, and 1000). (e,f) Sensitivity of $\phi_{xy}$ and $\phi_z$ to the interaction cutoff radius $r_c$, evaluated for 120 length-matched 7TM proteins ($375 < N \le 475$ residues, $s = 16$) with $r_c = 8, 10, 12$, and 15 Å.

### S2. ROBUSTNESS AND FINITE-SIZE SCALING OF DIRECTIONAL CORRELATIONS

#### S2.1 Robustness of directional correlation scaling

At $s = 16$, correlation functions from different topological classes show similar directional finite-size scaling after normalization by the corresponding molecular extents (Fig. S2 a,b). Within the size-controlled subset, the rescaled curves collapse separately for the in-plane and normal directions, supporting a common scaling behavior across transmembrane-protein topologies.

To test the numerical robustness of the extracted correlation lengths, we reconstructed the directional correlation functions using $K = 20, 50, 100, 200$, and 1000 nonzero modes. Increasing $K$ modifies the amplitudes and short-distance structure of the correlation profiles, whereas the zero-crossing positions and corresponding correlation lengths remain nearly unchanged (Fig. S2 c,d). The directional correlation lengths are therefore governed predominantly by the low-frequency collective spectrum.

We further varied the interaction cutoff radius $r_c$. Smaller cutoffs, particularly $r_c = 8$ Å, produce slightly shorter correlation lengths because of reduced network connectivity, whereas the correlation profiles progressively converge as $r_c$ increases and become nearly stable for $r_c \ge 10$ Å (Fig. S2 e,f). Together, these tests show that the directional correlation lengths are robust to both mode truncation and reasonable variations in network connectivity.

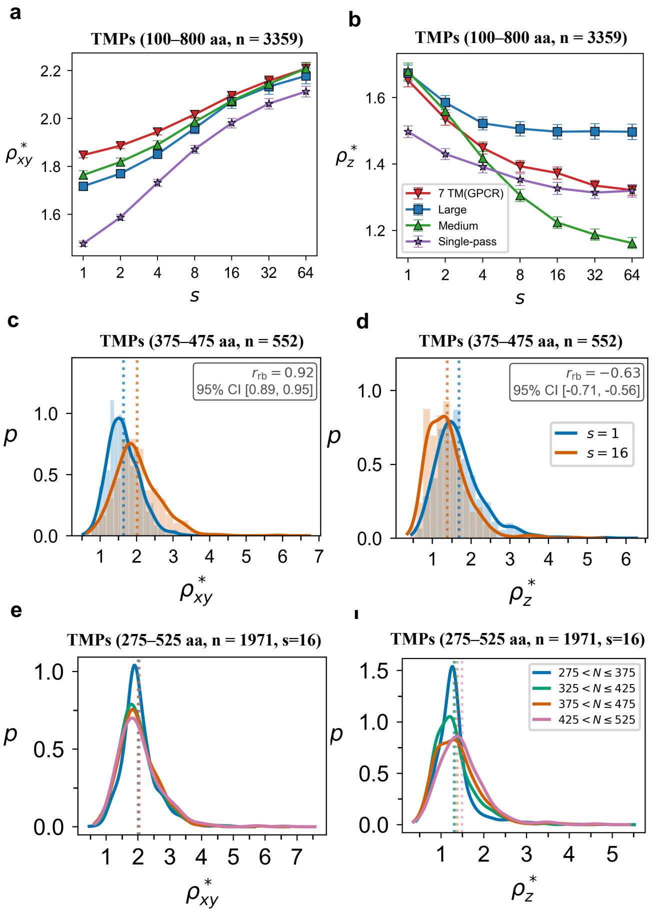


FIG. S3. Finite-size scaling and opposing shifts of directional correlation lengths under membrane anisotropy. (a,b) Reduced directional correlation lengths $\rho^*_{xy}$ and $\rho^*_z$ as functions of the stiffness ratio $s$ across the complete dataset of 3,359 transmembrane proteins. (c,d) Probability density distributions of $\rho^*_{xy}$ **(c)** and $\rho^*_z$ **(d)** for a size-controlled ensemble of 552 transmembrane proteins ($375 \leq N \leq 475$ residues) spanning four topological classes, comparing $s = 1$ and $s = 16$. (e,f) Probability density distributions of $\rho^*_{xy}$ **(e)** and $\rho^*_z$ **(f)** at $s = 16$ across four chain-length groups spanning 1,971 transmembrane proteins ($275 \leq N \leq 525$ residues), showing the weak residual dependence of the reduced correlation lengths on protein size.

#### S2.2 Finite-size scaling and opposing directional shifts under membrane anisotropy

Because transmembrane proteins span a broad range of molecular sizes and shapes, the dimensional correlation lengths $\xi_{xy}$ and $\xi_z$ naturally vary with the corresponding molecular dimensions. We therefore use the reduced correlation lengths $\rho^*_{xy} = \xi_{xy}/L_{xy}$ and $\rho^*_z = \xi_z/L_z$ to separate this finite-size scaling from the directional reorganization induced by membrane anisotropy. If the directional correlation lengths remain proportional to the corresponding molecular dimensions, $\rho^*_\alpha$ should be approximately independent of protein size.

Across the complete dataset of 3,359 transmembrane proteins, increasing the stiffness ratio produces opposite shifts in the two reduced correlation lengths: $\rho^*_{xy}$ increases monotonically with $s$, whereas $\rho^*_z$ decreases (Fig. S3 a,b). The direction of these shifts is consistent across all topological classes, although their magnitude remains topology dependent. Thus, membrane anisotropy systematically extends in-plane correlations while shortening normal correlations without introducing a fixed correlation length.

To assess the consistency of this directional reshaping at the level of individual proteins, we compared the distributions of $\rho^*_{xy}$ and $\rho^*_z$ between $s = 1$ and $s = 16$ for a size-controlled ensemble of 552 transmembrane proteins ($375 \leq N \leq 475$ residues) spanning four topological classes (Fig. S3 c,d). Matched-pairs rank-biserial correlations were calculated between the two stiffness conditions, with 95% confidence intervals estimated by percentile bootstrap using 5,000 resamples. The in-plane correlation length shifts systematically toward larger values, with $r_{\mathrm{rb}} = 0.92$ (95% CI: $[0.89, 0.95]$), whereas the normal correlation length shifts toward smaller values, with $r_{\mathrm{rb}} = -0.63$ (95% CI: $[-0.71, -0.56]$). These results confirm that the opposing directional shifts are not driven by ensemble averaging but occur consistently across individual proteins.

Additionally, we tested the residual dependence of the reduced correlation lengths on protein size. At $s = 16$, the distributions of $\rho^*_{xy}$ and $\rho^*_z$ were compared across four chain-length groups spanning 1,971 proteins within the 275–525 residue range (Fig. S3 e,f). The distributions and their mean values strongly overlap across size categories, showing that normalization by the corresponding directional molecular extents substantially reduces the dependence of the correlation lengths on protein size. Together with the directional collapse of the correlation functions, these results support a common finite-size scaling of long-range correlations while showing that membrane anisotropy reshapes this organization differently within the membrane plane and along its normal.

### S3. MECHANICAL ORIGIN OF THE OPPOSING DIRECTIONAL REORGANIZATION

The directional reorganization in Fig.3 originates from mechanical compatibility across the native contact network. The membrane constraint acts uniformly on all native contacts, but its effect depends on contact orientation. For tilted contacts, normal and lateral relative displacements enter the same bond extension, so collective motion must coordinate the two components across the network. Increasing the stiffness ratio changes these compatibility conditions and thereby reorganizes the collective-mode spectrum.

#### S3.1 Mechanical compatibility of tilted contacts

Let $\boldsymbol{q}_i$ denote the displacement of residue $i$, and let $\boldsymbol{e}_{ij} = (\boldsymbol{r}_i^0 - \boldsymbol{r}_j^0)/r_{ij}^0$ be the native unit vector of a contact between residues $i$ and $j$. We decompose the contact orientation and relative displacement as

$$\boldsymbol{e}_{ij} = \boldsymbol{e}_{ij}^{xy} + e_{ij}^z \hat{\boldsymbol{z}}, \qquad \Delta\boldsymbol{q}_{ij} = \Delta\boldsymbol{q}_{ij}^{xy} + \Delta q_{ij}^z \hat{\boldsymbol{z}}, \tag{S7}$$

where $\Delta\boldsymbol{q}_{ij} = \boldsymbol{q}_i - \boldsymbol{q}_j$.

With the membrane normal aligned with the $z$ axis, define

$$\mathsf{D}_s^{(3)} = \sqrt{s}\,\mathsf{P}_{xy} + \mathsf{P}_z = \mathrm{diag}(\sqrt{s}, \sqrt{s}, 1), \qquad \mathsf{D}_s = \mathbf{I}_N \otimes \mathsf{D}_s^{(3)}. \tag{S8}$$

The elastic energy of the anisotropic network is

$$U_s = \frac{\gamma}{2} \sum_{\substack{i<j \\ r_{ij}^0 < r_c}} \left[\sqrt{s}\,\boldsymbol{e}_{ij}^{xy} \cdot \Delta\boldsymbol{q}_{ij}^{xy} + e_{ij}^z \Delta q_{ij}^z\right]^2. \tag{S9}$$

For a tilted contact, the in-plane and normal displacements therefore contribute to the same extension. A low-strain deformation approximately satisfies

$$\sqrt{s}\,\boldsymbol{e}_{ij}^{xy} \cdot \Delta\boldsymbol{q}_{ij}^{xy} + e_{ij}^z \Delta q_{ij}^z \approx 0. \tag{S10}$$

Increasing $s$ rotates this locally compatible direction in displacement space. A collective mode must accommodate these rotated conditions simultaneously over many contacts with different orientations, so the membrane constraint reorganizes global motion even though the native coordinates and contact topology remain unchanged.

Equation S10 also explains why extended normal motion is generically mixed rather than purely normal. If the lateral component vanishes, the compatibility condition reduces to $e_{ij}^z \Delta q_{ij}^z \approx 0$, so tilted contacts with $e_{ij}^z \neq 0$ require neighboring residues to have nearly equal normal displacements. Across a connected contact network, this drives the motion toward a uniform $z$ translation, which does not contribute to internal fluctuations. Nontrivial long-range normal coordination therefore requires finite lateral displacements to accommodate variations in $z$ motion across differently oriented contacts. In membrane proteins, such mixed collective modes naturally include coordinated rocking, tilting, and repacking of transmembrane helices.

The network form of Eq. S9 is

$$\mathbf{H}_s = \mathsf{D}_s \mathbf{H}_1 \mathsf{D}_s. \tag{S11}$$

Thus, anisotropy changes the mechanical metric of the complete contact network rather than introducing an independent penalty on a selected set of contacts.

#### S3.2 Reweighting of the collective-mode spectrum

Let the non-rigid eigendecomposition of the isotropic Hessian be

$$\mathbf{H}_1 = \mathbf{V}\boldsymbol{\Lambda}\mathbf{V}^T. \tag{S12}$$

Defining $\mathbf{M}_s = \mathsf{D}_s\mathbf{V}$ and $\mathsf{G}_s = \mathbf{V}^T \mathsf{D}_s^2 \mathbf{V}$, the transformed modes are generally nonorthogonal and the exact covariance becomes

$$\mathbf{C}_s = k_{\mathrm{B}}T\,\mathbf{M}_s \mathsf{G}_s^{-1} \boldsymbol{\Lambda}^{-1} \mathsf{G}_s^{-1} \mathbf{M}_s^T. \tag{S13}$$

For isotropic mode $\boldsymbol{v}_a$, define its in-plane participation as

$$p_a = \boldsymbol{v}_a^T (\mathbf{I}_N \otimes \mathsf{P}_{xy})\, \boldsymbol{v}_a, \qquad 0 \le p_a \le 1. \tag{S14}$$

The corresponding diagonal element of $\mathsf{G}_s$ is $1 + (s-1)p_a$. Neglecting off-diagonal mode mixing only to expose the physical trend, the modal contributions to the two directional covariances, apart from direction-wide factors that cancel upon normalization, scale as

$$\widetilde{u}_a^{xy} \propto \frac{p_a}{\lambda_a[1+(s-1)p_a]^2}, \qquad \widetilde{u}_a^{z} \propto \frac{1-p_a}{\lambda_a[1+(s-1)p_a]^2}. \tag{S15}$$

All numerical calculations retain the full anisotropic Hessian and complete mode mixing.

For $s \gg 1$ and $p_a \gg s^{-1}$, the common factor $s^{-2}$ cancels within each normalized directional correlation function, leaving

$$\widetilde{u}_a^{xy} \propto \frac{1}{\lambda_a p_a}, \qquad \widetilde{u}_a^{z} \propto \frac{1-p_a}{\lambda_a p_a^2}. \tag{S16}$$

The same anisotropic filter therefore acts through different directional projections. At large $s$, the normal covariance becomes more strongly biased toward modes with small in-plane participation than the in-plane covariance. Because nontrivial long-range normal coordination requires mixed modes with finite $p_a$, this redistribution removes the modes that connect distant regions along $z$, fragmenting $\phi_{ij}^z$ and shortening $\rho_z^*$.

For the in-plane component, the overall reduction in fluctuation amplitude is removed by normalization. In the networks studied here, the surviving soft spectrum is increasingly dominated by spatially coherent motion whose in-plane projection extends across multiple helices and structural regions. The resulting merging of $\phi_{ij}^{xy}$ domains increases $\rho_{xy}^*$. The opposing directional trends therefore reflect two projections of the same reorganization of mechanical compatibility.

### S4. DIRECTIONAL CORRELATIONS IN OLIGOMERIC PROTEIN ASSEMBLIES

To assess whether the directional-correlation framework extends beyond the monomeric structures analyzed in the main text, we examined the $\mu$-opioid receptor dimer (PDB ID: 4DKL) as a representative GPCR assembly (Fig. S4a). As shown in Fig. S4b,c, the isolated single chain exhibits zero crossings at $\xi_{xy} = 20.9$ Å and $\xi_z = 16.4$ Å, whereas the assembled dimer yields $\xi_{xy} = 24.0$ Å and $\xi_z = 20.7$ Å. Both correlation lengths

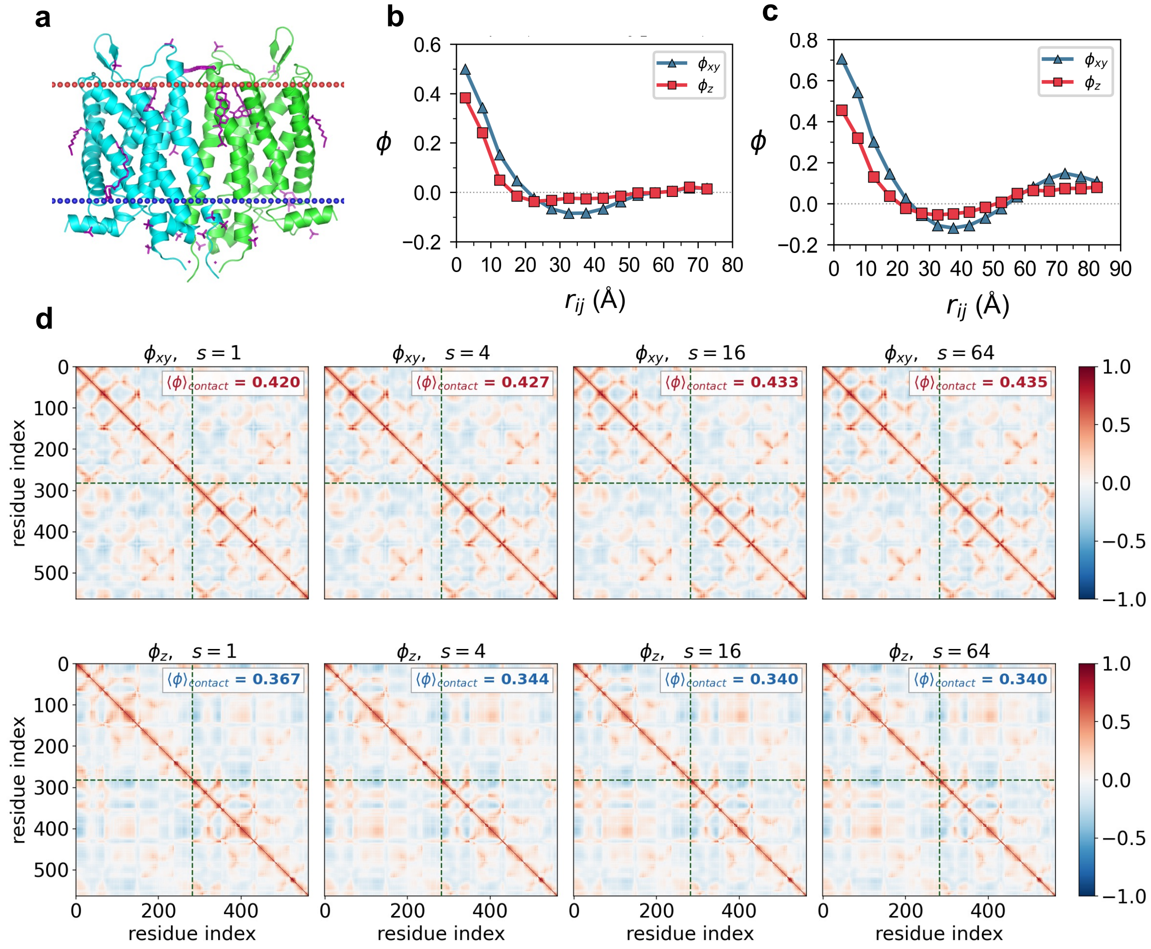


FIG. S4. Directional correlations in a dimeric membrane-protein assembly. (a) Structural model of the GPCR $\mu$-opioid receptor dimer (PDB ID: 4DKL; membrane-embedded model retrieved from the OPM database, Main Text Ref. 55). (b,c) Directional correlation functions $\phi_{xy}$ (blue triangles) and $\phi_z$ (red squares) as functions of the three-dimensional native inter-residue distance $r_{ij}$ at $s = 16$, shown for the isolated single chain **(b)** and the assembled dimer complex **(c)**. (d) Directional correlation matrices of the assembled $\mu$-opioid receptor dimer under membrane constraints. $\phi_{ij}^{xy}$ (top) and $\phi_{ij}^{z}$ (bottom) are shown for $s = 1$, 4, 16, and 64. Dashed lines separate the two protomers; $\langle\phi_{ij}^{\alpha}\rangle_{\text{contact}}$ denotes the mean correlation over native contacts ($r_{ij} < 10\,\text{Å}$).

increase upon assembly, consistent with the larger spatial extent of the dimer, while the directional separation $\xi_{xy} > \xi_z$ is preserved. Because the chain and dimer share the same membrane reference frame and anisotropic constraint, the comparison isolates the additional effect of inter-subunit coupling.

We further examined residue-level correlations as membrane anisotropy increased in the assembled dimer (Fig. S4d). The mean native-contact correlation within the membrane plane, $\langle\phi_{ij}^{xy}\rangle_{\text{contact}}$, increases from 0.420 at $s = 1$ to 0.433 at $s = 16$ and 0.435 at $s = 64$, whereas the normal component, $\langle\phi_{ij}^{z}\rangle_{\text{contact}}$, decreases from 0.367 at $s = 1$ to 0.340 at $s = 16$ and remains suppressed at $s = 64$. The complex therefore shows the same opposing directional trend as the monomeric setup, with increasing anisotropy enhancing in-plane contact correlations while reducing those along the membrane normal.

### S5. SUPPLEMENTARY DATA AND CODE AVAILABILITY

**Supplementary Data:** The curated mutagenesis benchmark dataset, comprising 1,452 experimentally characterized transmembrane mutation positions across 20 human GPCRs compiled from GPCRdb, is deposited on Zenodo (DOI: 10.5281/zenodo.21993854).

**Code Availability:** Code used for the analyses, including imANM simulations, directional decomposition and rdDFI calculations, and GPCR benchmark evaluation, is publicly available on GitHub (`https://github.com/Cosmoto-jian/rdDFI`).
